# Dynamical properties of self-sustained and driven neural networks

**DOI:** 10.1101/2023.07.15.549166

**Authors:** Jules Bouté, Alain Destexhe

## Abstract

In the awake brain, cerebral cortex displays asynchronous-irregular (AI) states, where neurons fire irregularly and with low correlation. Neural networks can display AI states that are self-sustained through recurrent connections, or in some cases, need an external input to sustain activity. In this paper, we aim at comparing these two dynamics and their consequences on responsiveness. We first show that the first Lyapunov exponent (FLE) can differ between self-sustained and driven networks, the former displaying a higher FLE than the late. Next, we show that this impact the dynamics of the system, leading to a tendency for self-sustained networks to be more responsive, both properties that can also be captured by mean-field models. We conclude that there is a dynamical and excitability difference between the two types of networks besides their apparent similar collective firing. The model predicts that calculating FLE from population activities in experimental data could provide a way to identify if real neural networks are self-sustained or driven.

## 1 Introduction

Neural networks are a core subject of analysis when it comes to model the brain. They can behave in vastly different ways, and we will focus here on the so-called Asynchronous Irregular (AI) state as defined (with the other types of behavior) in (Brunel, 2000) and first observed in neural networks in (Amit & Brunel, 1997; van Vreeswijk & Sompolinsky, 1996; Vreeswijk & Sompolinsky, 1998). This means neurons fire without any significant coherence between each other and that single single neurons would not have a regular pattern of activity.

That type of activity is similar to that observed in awake animals (Matsumura, Cope, & Fetz, 1988; Steriade, Timofeev, & Grenier, 2001; Destexhe, Rudolph, & Paré, 2003; Lee, Manns, Sakmann, & Brecht, 2006), including primates and humans (Dehghani et al., 2016) and therefore represent a basic type of activity of prime importance. This is why it is useful for networks to be able to reproduce it, and how it is an interesting approach when testing common models.

Different models of single neuron and networks allow to reproduce these dynamics, such as the AdEx model (Brette & Gerstner, 2005; Naud, Marcille, Clopath, & Gerstner, 2008), where it was shown that AdEx networks can display AI states (Destexhe, 2009), which will be the focus of this study. The well-known Hodgkin-Huxley model (Hodgkin & Huxley, 1952), which can also display AI states (Carlu et al., 2019), which we use here for comparison.

While the global activity can be reproduced well with such models, they can also have a large range of different behaviors as shown for example in (Naud et al., 2008), and a change of parameters within the single neurons models or the network model can lead to vastly different dynamics. Although some of those dynamics can be obvious to differentiate (such as a change in the average firing rate that one would adapt to reproduce specific brain data, or reactions to specific inputs that could also be modulated), there are other sets of parameters that would appear very similar.

A good example is the existence of two different types of networks : Driven networks, that need an external drive such as in (Brunel, 2000; Zerlaut, Chemla, Chavane, & Destexhe, 2017) and Self-sustained networks, that just need to be started to have an activity such as in (Vogels & Abbott, 2005; Destexhe, 2009).

Those two types of networks appear very similar if we overlook the need for an external drive, and the two of them could even be use with an external drive. Therefore, if they appear so similar, would those two types of networks behave the same way ? Can those two categories be meaningful and predictive of different dynamics altogether ? This are the questions we will try to answer in this study.

To do so, we use tools from dynamical systems (Eckmann & Ruelle, 1985; Ruelle, 1989) applied to computational neuroscience (Faure & Korn, 2001; Izhikevich, 2007; Cessac, 2009), and in particular the Lyapunov Exponents (Cessac, 2009; Ruelle, 2009) (as defined in (Wolf, Swift, Swinney, & Vastano, 1985; Eckmann, Kamphorst, Ruelle, & Ciliberto, 1987)), which we apply here to the two types of networks. We further probe the networks by studying their responsiveness to external synaptic inputs, as in (Yger, El-Boustani, Destexhe, & Frégnac, 2011; Zerlaut et al., 2017), allowing us to see how such network states could be used to process or transmit information in the brain (Zerlaut & Destexhe, 2017)or generate pathological activity such as epilepsy (Depannemaecker, Carlu, Bouté, & Destexhe, 2022).

Finally, we will compare our findings with mean-field models of the two network types, following (El-Boustani & Destexhe, 2009; Zerlaut et al., 2017; di Volo, Romagnoni, Capone, & Destexhe, 2019), allowing us to see if the concepts and properties of Driven and Self-sustained networks are equivalent to that of mean-field models.

## 2 Methods

### 2.1 Neural network model

We now present the model of single neurons that we use, and how said single neurons are linked together to form a network.

It is useful to point that we took parameters such that our networks would be Asynchronous Irregular, as defined in the Introduction.

#### 2.1.1 AdEx model

To simulate a neural network, we fist need to simulate single neurons. We decided to focus on the Adaptative Exponential Integrate and Fire model (Brette & Gerstner, 2005; Naud et al., 2008), or “AdEx” model here. In the following, unless otherwise stated, we focus by default on the AdEx network.

Each neuron in the AdEx network is described by Eq.(1) and Eq.(2) as follows:

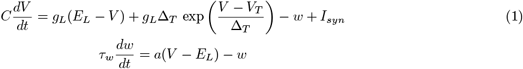

When the membrane potential crosses a threshold, a spike is emitted, and the system is reset:

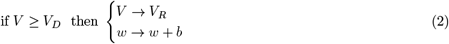

Then, to create a biological networks, we simulate two kind of neurons, that we call populations : excitatory and inhibitory neurons. As the name suggests, when the first kind of neuron will spike, it will increase the chance of spiking of other neurons, while for the second kind it will decrease that chance. We simulate a total of 10 000 neurons, 2000 being inhibitory - that we will call FS for Fast Spiking neurons - and 8000 being excitatory - that we will call RS for Regular Spiking neurons.

In order to influence other neurons, they need to be connected to each other. We choose a random Erdős–Rényi one way connection of 5%, so every neurons is on average connected to 400 excitatory neurons and 100 inhibitory neurons, and receives input for the same amount of neurons.

On top of those neurons, we will often use an external excitatory input from 8000 excitatory Poisson “neurons”. The corresponding excitatory input will be used as a drive (see 3.1) or as a perturbation (see Section 3.3). The strength of that drive/perturbation will be given in term of the average firing rate of the Poisson neurons (for example, 2Hz).

#### 2.1.2 Hodgkin and Huxley model

Although the primary model we used is the AdEx model, we also studied the Hodgkin Huxley (Hodgkin & Huxley, 1952) model, that we will call “HH” model.

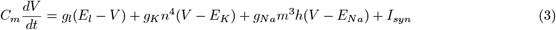

with gating variables (in ms):

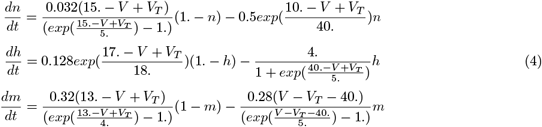

Apart from the single neuron model, HH networks function the same way as AdEx networks described previously.

### 2.2 Mean-Field

We used mean-field models of AdEx networks, using the model defined in (di Volo et al., 2019). This mean-field model was originally based on a Master Equation formalism developed for balanced networks of integrate-and-fire neurons (El-Boustani & Destexhe, 2009). This model was first adapted to AdEx networks of RS and FS neurons (Zerlaut et al., 2017), and later modified to include adaptation (di Volo et al., 2019). This latter version corresponds to the following equations (using Einstein’s index summation convention where sum signs are omitted and repeated indices are summed over):

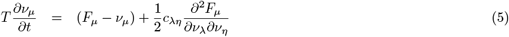

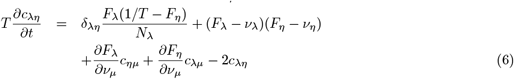

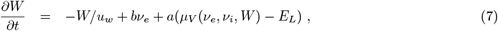

where *μ* = {*e, i*} is the population index (excitatory or inhibitory), *ν*_*μ*_ the population firing rate and *c*_*λη*_ the covariance between populations *λ* and *η. W* is a population adaptation variable divolo2019. The function *F*_*μ*={*e,i*}_ = *F*_*μ*={*e,i*}_(*ν*_*e*_, *ν*_*i*_, *W*) is the transfer function which describes the firing rate of population *μ* as a function of excitatory and inhibitory inputs (with rates *ν*_*e*_ and *ν*_*i*_) and adaptation level *W*. These functions were estimated previously for RS and FS cells and in the presence of adaptation (di Volo et al., 2019).

At the first order, i.e. neglecting the dynamics of the covariance terms *c*_*λη*_, this model reduces to:

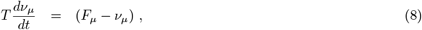

together with Eq. 7. This system is equivalent to the well-known Wilson-Cowan model (Wilson & Cowan, 1972), with the specificity that the functions *F* need to be obtained according to the specific single neuron model under consideration. These functions were obtained previously for AdEx models of RS and FS cells (Zerlaut et al., 2017; di Volo et al., 2019) and the same are used here.

### 2.3 Lyapunov exponent algorithm

Considering the way we simulated the networks and mean-field, with differential equations, and the apparent differences in term of activity due to subtle differences, we chose to analyse the simulations with a dynamical systems lens, meaning we will study the trajectory of the dynamics on the phase space : the space of the dynamic of the different variables. To do so, we mainly focus on one of the primary tools of dynamical system analysis : Lyapunov exponents, that we will hereafter write LE.

LE are often (Eckmann & Ruelle, 1985; Pikovsky & Politi, 2016) defined as a way to observe the stability of specific points of the system : fixed points, points in which the differential equations goes to 0, inducing an absence of evolution of the trajectory. Positive exponent means the point is unstable : a small perturbation will just grow and goes away from the fixed point. Negative exponents mean the point is stable : small perturbation will just decrease and come back to the fixed point. In multidimensional systems, there is one LE per dimension, meaning that, when look at the phase space, there could be a mixture of stable and unstable directions from a fixed point.

While important, fixed point analysis is not the only thing that can be done with LE. A generalized version of this analysis allows to compute the stability of a whole system (Pikovsky & Politi, 2016) (or at least of a specific attracting object withing it), which is what interested us. Here, we would considered system that are bounded, meaning that if there is some unstable directions due to positive exponents, they would not go to infinite : the perturbation would just grow, until the two path of the original and perturbed trajectory are completely uncorrelated. This is called deterministic chaos (Strogatz, 2019). Chaotic systems have, on appearance, similar properties to random, noisy ones, but they are entirely deterministic which allows for a whole different kind of analysis and predictive power. Knowing if a system is chaotic or not, is an important outcome of the estimation of LE, as at least one positive exponent is a defining feature of deterministic chaos (Wolf et al., 1985; Eckmann et al., 1987). It was also shown that many systems in the brain have properties consistent with chaotic systems (Korn & Faure, 2003).

Finding LE requires different algorithms, depending on the situation, from simple to complex one, especially when the goal is to find the complete spectrum and the associated direction (Pikovsky & Politi, 2016). When the differential equations are available, the basic idea behind the algorithm is to do some perturbation, let the system evolve, and check if the distance between the original and the perturbed point has grown or shrunk. The logarithm of this distance will be the exponent. Of course, this method should be done a lot of time to converge to a good value, as there will be some differences in different regions of the system. As can be guessed, this only allows to compute the largest LE, that we will call FLE for “First Lyapunov Exponent”. We will rank each LE from higher to lower. If one wants to obtain the rest of the spectrum, it is usually requires to go to an orthogonal subspace from the direction of the first exponent, in order to cancel its effect, and to start over again for each exponent. For more detail, see (Pikovsky & Politi, 2016), as finding more than the FLE is outside the scope of this work.

We said previously that understanding if a system was chaotic or not was of prime importance, which is why this kind of analysis was not restricted to analysing systems for which the equations were known : algorithm were created to compute the LE (and especially the FLE, as it was the required one to determine if a system is consistent with chaos) from time series (Wolf et al., 1985; Eckmann et al., 1987), in order to apply it to real data.

Those algorithms work basically the same way as what was explained before, except that nearest neighbors are used instead of small perturbation. From then, the trajectory of the original point and the neighbor are observed for a certain amount of time steps in order to compute the change of distance between them. Finding the right amount is important : too many, and if the system is chaotic, all correlation between the two trajectory will be lost, meaning the distance would be meaningless. Too short, on the other hand, would not leave enough time for the trajectory to evolve, and would result in an increasingly longer computational time, on top of adding more potential errors. This is why we always tested two time steps, written as “Dt”, to ensure the effect we found would be stable within a certain range. As before, we would take the logarithm of that distance that would have either shrunk or grow, to compute the FLE. It is again required to repeat the process and do an average on many times to converge to a meaningful value. The complete algorithm that was used is given in the supplementary section.

It is important to note that those algorithm works very well to find positive LE (not necessarily only the first one), but suffer from important bias when they try to compute negative ones (Wolf et al., 1985; Eckmann et al., 1987). In our case, only the FLE was positive, and the second one would be either 0 or negative, which is why we stopped our analysis in computing only the FLE of the systems (or the whole spectrum when fixed points where considered, in the case of the mean-field, as it would only be an analytical computation).

## 3 Results

We wanted to analyse the system using all variables, but it proved to be impossible due to their sheer number, as all tested algorithms relied on a distance metric (generally to find nearest neighbors) that always broke due to the curse of dimensionality.

Therefore, we needed fewer variables, which is why we chose variables that were the most relevant for the system, on top of being the ones used to simulate a mean-field of said system, allowing for a great connection between two different representations of the same phenomenon. Two obvious choices were the average firing rates of the excitatory and inhibitory population, which gives an idea of the average behavior of the system, and the last one is the average adaptation variable, only present in the excitatory population, which is a specificity of the AdEx model, and therefore an important feature to represent. For HH, we will choose the other 6 variables : n,h and m for of the inhibitory and excitatory populations.

It is worth noting that the first two variables are of a different kind from the last ones : the adaptation (or mn,h and m) is an intrinsic part of the description of the system, having its own dynamical equations, while the firing rates are product of the system, when the membrane potential reaches a threshold.

We start by describing the self-sustained and driven types of networks, then we compute their Lyapunov exponents, and next their responsiveness to external inputs.

### 3.1 Driven and self-sustained networks

Neural networks can vary in term of the model of single neuron or the type of connectivity as shown previously, but we want to add another difference : what we call the type of networks, which can be driven or self-sustained.

#### 3.1.1 AdEx

Driven networks are the networks that require an external (here Poissonian) input to produce an activity (see Fig.1(A-B)). As most parameters will work to create driven networks, they are also the easiest to obtain (if we allow an external drive).

**Figure 1:**
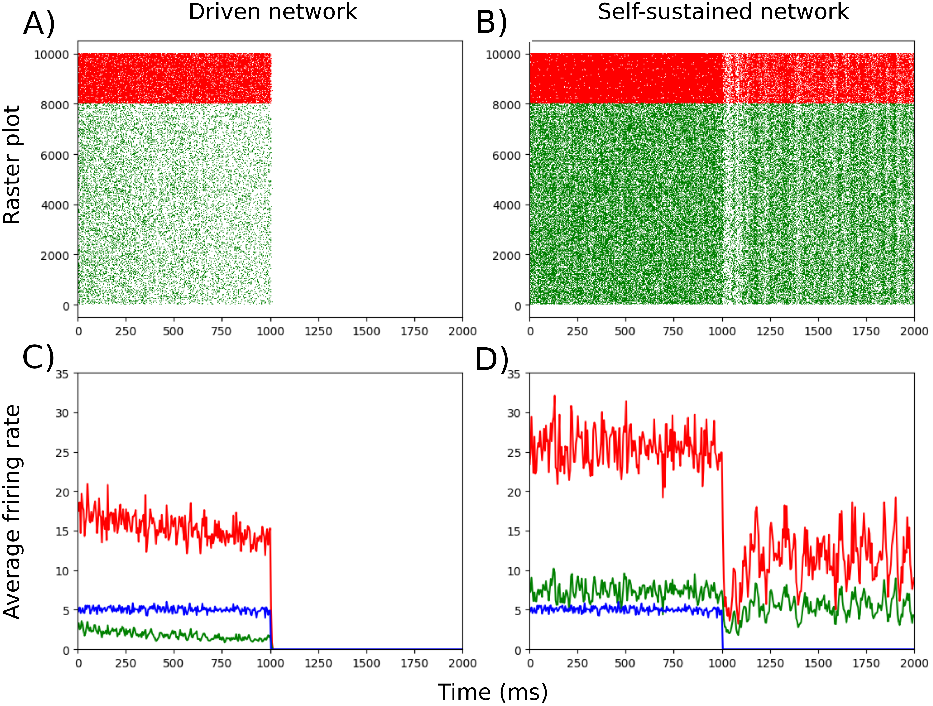
Illustration, for an AdEx network, of driven networks (A) and B)) and self sustained networks (C) and D)) with an external Poisson drive that stops at 1000ms. A) and C) represent raster plots, with green being RS neurons and red being FS neurons. B) and D) represent the average firing rate, with the same colour code and blue being the Poisson input. As can be seen, when said Poisson input stops, the driven networks immediately stop any activity, while the self-sustained reduce but maintain its activity.

Self-sustained networks on the other hand only needs an initial “kick” to start, and will, as the name suggest, have a self-sustained activity after (see (see Fig.1(C-D)).

Both types of networks are interesting for different reasons : driven networks might be more realistic, as a small network is never cut out of the rest of the brain, and has no specific reason to sustained itself. On the other hand, self-sustained networks allows for an easier analysis of their dynamics, as there is no external input and most of all no sources or randomness (through the poissonnian input) that will affect them. Self-sustained are more difficult to obtained than driven : on top of requiring some fine tuning (although an important range of parameters is compatible with it), it also requires specific noise realizations (so a specific random connectivity and a specific kick), as many noise realizations will just die quickly (in less than a second, with a kick of 100ms).

It is important to note that, while some change of parameters and study of the dynamics of the system gives good insight on how to obtain a self-sustained network, it is beyond the scope of this study to prove they were actually self-sustained forever and it was only checked that they would have a seemingly stable activity for longer than the typical length of simulation we needed (more than 20s), as even if it is only a transient phase, its lifetime would be big considerin the size of the network (Tél & Lai, 2008). While the self-sustained networks do not require it, we sometimes add a drive to them for the sake of comparison with the driven system.

#### 3.1.2 HH

We obtained the same two types of networks for HH networks, as can be seen if Fig.2. Here we can see that the firing rates look very similar to the AdEx networks, albeit being a bit higher due to a choice of parameters, especially for the self-sustained HH network (while it could have been possible, in theory, to have lower firing rates, finding the correct parameters to obtain a self-sustained activity is a difficult task, so those parameters were kept).

**Figure 2:**
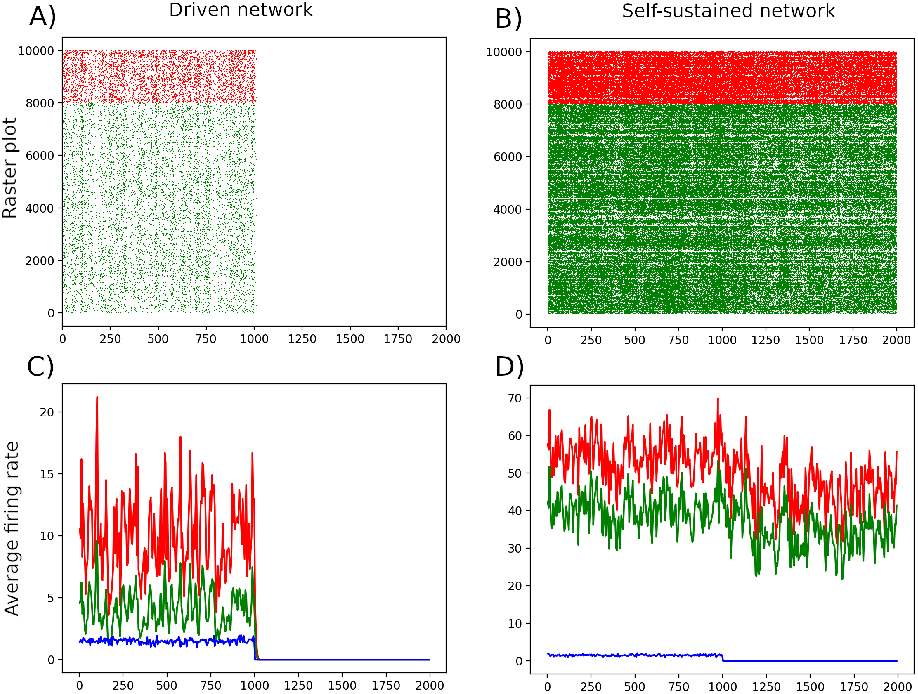
Illustration, for an HH network, of driven networks (A) and B)) and self sustained networks (C) and D)) with an external Poisson drive that stops at 1000ms. A) and C) represent raster plots, with green being RS neurons and red being FS neurons. B) and D) represent the average firing rate, with the same colour code and blue being the Poisson input. As can be seen, when said Poisson input stops, the driven networks immediately stop any activity, while the self-sustained reduce but maintain its activity.

#### 3.1.3 Mean-field

With the mean-field, it was also possible to obtain Driven like and Self-sustained like activity, as can be seen in Fig.3 in the sense that with some parameters, the activity would go to 0 without external drive for some networks, while it would go to some non-zero values for some other. Due to the very different dynamical nature of the mean-field, as it has attractive fixed points instead of chaos, as we will see, it is hard to know if the Driven and Self-sustained networks are of the same type as the corresponding networks.

**Figure 3:**
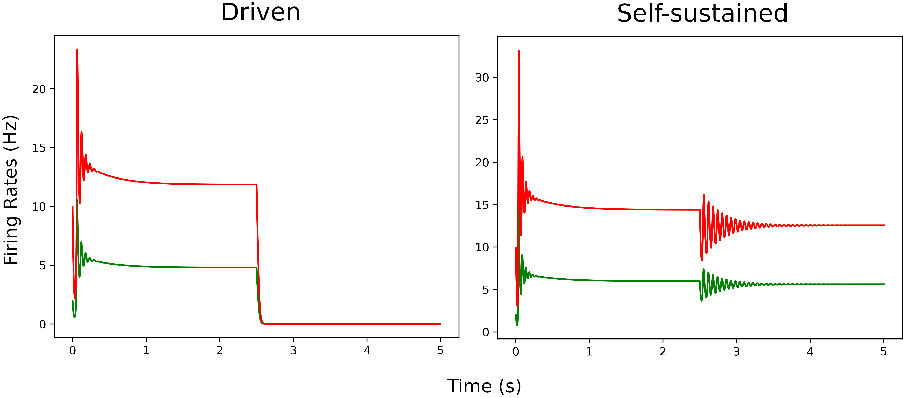
Driven (left) and self-sustained (right) mean-field, with fixed external input at 1.5Hz for 2.5s and then no input for the last 2.5s.

We can see some oscillations after a change of state (at the beginning for each network simulation, and when we stop the external input for the Self-Sustained network). This is most likely an artefact, due to the mean-field being created to simulate steady states, and not particularly the transient state when there is a change. This is why we will only consider the steady states values.

### 3.2 Lyapunov exponents comparison between Driven and Self-sustained systems

Using the algorithm described earlier, we computed the FLE on a dynamical system made from the 3 dimensions described previously (8 for HH). Usually, the algorithm requires to obtain the phase space using Takens reconstruction, but here we force it to operate in the three dimensions we constructed, as they 1) make sense to represent all the system (the normal algorithm often assume we only have access to partial information) and 2) make it easier to identify with the mean-fields (for AdEx as the mean-field analysis was only performed for it, although the same would have been true for an HH mean-field).

#### 3.2.1 AdEx

Fig.4 is a good way to get a representation of how the algorithm works, and what it means to have a given LE : we can see it is really a converging value of the whole system, that varies from time to time (or more accurately, depending on when we take it in the system) but still converges, which is why there is some meaning in talking about the FLE of the different systems.

**Figure 4:**
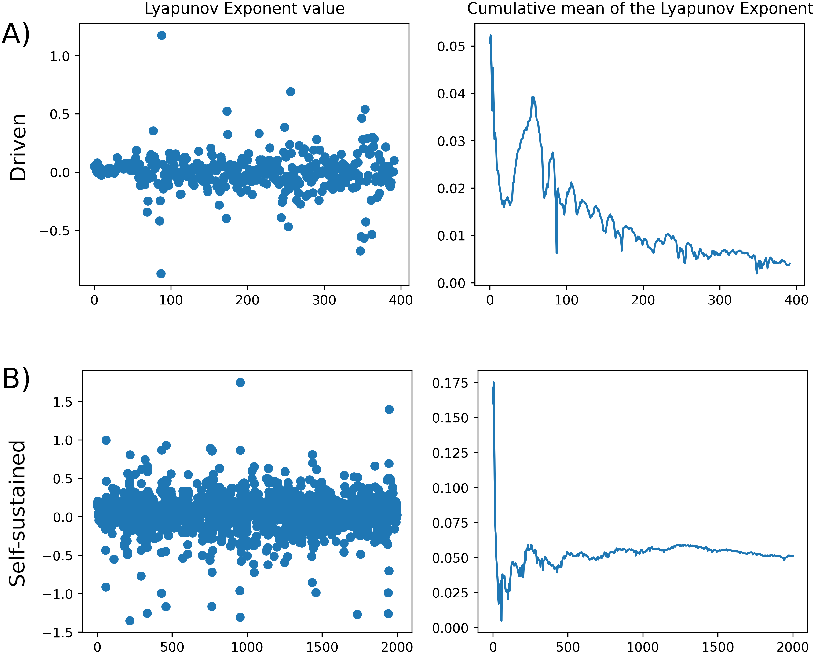
Example of FLE algorithm for typical self-sustained (up) and driven (down) AdEx networks. On the left, we have the values of the exponents at various steps. On the right, we have the cumulative mean. The self-sustained converges to 0.05 and the driven converges to 0.002

To explore the differences between Driven and Self-sustained networks, we computed the FLE for different conditions in Table.1

**Table 1:** Values of the FLE for the different conditions. There are two parts : the first one is computed with the two mean firing rates (FR) and the mean adaptation, while the second part is computed with the two mean membrane potentials (Vm) and the mean adaptation. The column represent different networks, 2 driven networks with different parameters, and two similarly different self-sustained networks. The lines represent 2 different time steps used in the algorithm, to ensure some robustness. Finally, each condition was repeated with a different noise realization (including both the drive and the connections in the network).

| FR | Driven 1 | Driven 2 | Self-Sustained 1 | Self-Sustained 2 |
| --- | --- | --- | --- | --- |
| Dt=10 | 0.01/0.003 | 0.007/0.0004 | 0.05/0.04 | 0.05/0.04 |
| Dt=20 | 0.004/0.003 | 0.006/0.002 | 0.04/0.04 | 0.05/0.07 |
| Vm |  |  |  |  |
| Dt=10 | 0.35/0.36 | 0.56/0.54 | 0.38/0.08 | 0.46/0.42 |
| Dt=20 | 0.35/0.31 | 0.41/0.41 | 0.20/0.21 | 0.33/0.32 |

Here we can see that while the FLE is always fairly weak, it is also positive (meaning we have chaotic systems) and an order of magnitude higher when we compare Driven to Self-sustained networks.

Or at least, that is what happens when we compute it with the firing rates. As can be seen in the 2^*nd*^ part of Table.1, when we use the intrinsic variables of the membrane potential Vm, the result is very different, showing no significant differences between Driven and Self-sustained networks. Interestingly, it also shows much higher LE. This shows that depending on the way we compute them, the exponents can be very different, and encourage us to use them more as a method of comparing than with their absolute values.

Table.2 shows similar results for a self-sustained network with an external drive, in order to see if the drive was the cause of the difference. As can be seen, results are similar to those for the “normal” self-sustained networks.

**Table 2:** Values of the FLE for the different conditions. The column represent different self-sustained networks with different drives, 2 1.5Hz drives and 2 5Hz drives. The lines represent 2 different time steps used in the algorithm, to ensure some robustness. Finally, each condition was repeated with a different noise realization (including both the drive and the connections in the network).

|  | 1.5Hz 1 | 1.5Hz2 | 5Hz 1 | 5Hz 2 |
| --- | --- | --- | --- | --- |
| Dt=10 | 0.05/0.05 | 0.06/0.06 | 0.06/0.05 | 0.06/0.06 |
| Dt=20 | 0.05/0.03 | 0.06/0.06 | 0.06/0.04 | 0.06/0.04 |

#### 3.2.2 HH

We also wanted to obtain the FLE for the HH networks, and as can be seen in Table.3 there is no difference between driven and self-sustained, which is why we did not investigate it further, focusing on the specificity of AdEx.

**Table 3:** Values of the FLE for the different conditions for the HH model. There are two parts : the first one is computed with the two mean firing rates (FR) and the mean adaptation, while the second part is computed with the two mean membrane potentials (Vm) and the mean adaptation. The column represent different networks, 1 driven and 1 self-sustained. The lines represent 2 different time steps used in the algorithm, to ensure some robustness. Finally, each condition was repeated with a different noise realization (including both the drive and the connections in the network).

| FR | Driven | Self-Sustained |
| --- | --- | --- |
| Dt=10 | 0.37/0.41 | 0.39/0.36 |
| Dt=20 | 0.26/0.27 | 0.31/0.26 |
| V<sub>m</sub> |  |  |
| Dt=10 | 0.49/0.51 | 0.48/0.43 |
| Dt=20 | 0.34/0.36 | 0.37/0.33 |

#### 3.2.3 Mean-Field

Finally, we also computed the LEs for the AdEx mean-field. This analysis is a bit different, as we actually only have three equations and everything is continuous. It was therefor possible to compute the entire spectrum directly using the torch library in python (torch.autograd.functional.jacobian). The results for a Driven and a Self-sustained networks, with and without external drives, are given in Table.4. We can see that we indeed have an attractive fixed points in all cases, as all exponents are negative. On top of that, the 1^*st*^ and 3^*rd*^ are, interestingly, always the same. The only difference arise for the 2^*nd*^ exponents, which is always higher for self-sustained, although only the case without external input seems to be significantly different.

**Table 4:** Values of the FLE for the different conditions for the AdEx mean-field. There are two parts : the first one is for a driven network, and the second one for a self-sustained network. The column represent each LE. The lines represent different amount of external inputs (in Hz)

| Driven | 1<sup>st</sup> exponent | 2<sup>nd</sup> exponent | 3<sup>rd</sup> exponent |
| --- | --- | --- | --- |
| Input = 0Hz | -2.00 | -66.67 | -66.67 |
| Input = 1.5Hz | -2.00 | -62.78 | -66.67 |
| Input = 3Hz | -2.00 | -65.96 | -66.67 |
| Self-sustained |  |  |  |
| Input = 0Hz | -2.00 | -44.18 | -66.67 |
| Input = 1.5Hz | -2.00 | -62.15 | -66.67 |
| Input = 3Hz | -2.00 | -65.77 | -66.67 |

### 3.3 Responsiveness

Finally, we wanted to see if the difference in the FLE (2^*nd*^ for the mean-field) would lead to differences in the response to an short external input between driven and self-sustained networks (mean-fields). As only AdEx showed differences, we only studied it for both the network and the mean-field.

To do so, we added an external poisson input, in blue in Fig.5. That added external input was done on top of the normal external drive for driven networks, and not in place of it, which is why we can see a non zero value everywhere that rise higher in the left of Fig.5 compared to the right where there is no external input apart from the perturbation.

**Figure 5:**
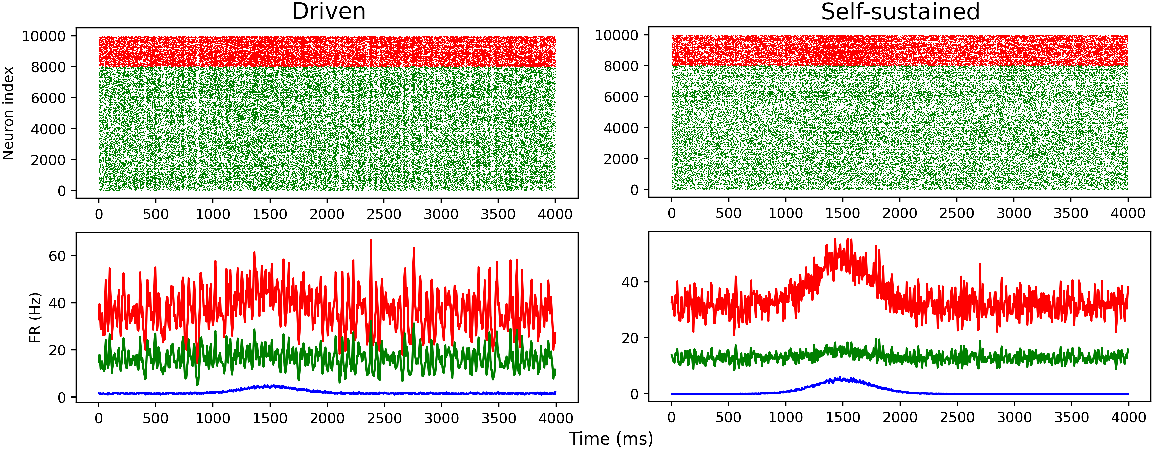
Response to a perturbation of 3Hz centered at 1500ms. Green represent the excitatory population, red is in inhibitory one, and blue is the external poisson drive. Up : raster plot. Down, average firing rates.Left part is for a driven network, right part for a self-sustained one.

We computed the response as follow :

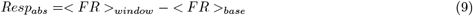

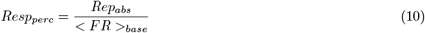

There are two measures of responsiveness that we used here : the first one is the absolute response, *Resp*_*abs*_, and is the difference between the average FR during a specific window chosen to be inside of the perturbation times, on one hand, and the average FR outside of the perturbation : the base FR. While interesting by itself and enough to compare, said, similar networks (mean-fields) with different external perturbation strength, we needed to compare networks (mean-fields) that had different base FR, and it could be that it would influence the absolute value of the response. To avoid that issue, we also computed the percentage response *Resp*_*perc*_, which is the *Resp*_*abs*_ divided by the mean base FR. Those two measures will be shown in the next Tables.

#### 3.3.1 Network

Fig.5 shows Driven and Self-sustained networks responding to an external perturbation of 3Hz. The responsiveness *Resp*_*abs*_ and the *Resp*_*perc*_, as defined previously, were reported (for different networks and different perturbation strength) in Table.5. Here we can see that Self-sustained networks seem to respond more than Driven ones, although the difference is not big an variability within Driven or Self-sustained networks is already high. The results are not as clear as they were for the FLE, but there seem to be a clear tendency.

**Table 5:** Response of the network with different external currents (in lines) for 2 Driven and 2 Self-sustained networks and for the two neuronal populations (in columns) : excitatory and inhibitory. In each cells, we have two *Resp*_*abs*_ for two noise realizations, followed by two *Resp*_*perc*_, also for two noise realizations.

| Driven | Exc. FR 1 | Exc. FR 2 | Inh. FR 1 | Inh. FR 2 |
| --- | --- | --- | --- | --- |
| Input = 3Hz | 0.67/0.84,<br>6.1%/7.5% | 0.11/0.11,<br>1.1%/1.2% | 3.28/3.58,<br>12.4%/13.4% | 2.22/2.26,<br>9.8%/9.8% |
| Input = 5Hz | 1.13/1.23,<br>10.2%/10.9% | 0.17/0.17,<br>1.9%/1.9% | 5.39/5.60,<br>20.3%/20.8% | 3.65/3.71,<br>16%/16% |
| Self-sustained |  |  |  |  |
| Input = 3Hz | 8.29/8.25,<br>25.7%/24.8% | 11.87/11.12,<br>32.1%/34.4% | 1.27/1.28,<br>9.9%/9.5% | 2.79/2.35,<br>19.5%/19.4% |
| Input = 5Hz | 13.12/12.84,<br>40.3%/38.2% | 17.91/17.65,<br>47.5%/53.2% | 1.83/1.80,<br>14.2%/13.3% | 4.05/3.66,<br>27.9%/29.7% |

#### 3.3.2 Mean-field

Fig.6 shows Driven and Self-sustained mean-field responding to an external perturbation of 3Hz. Contrary to Fig.5, there is no noise here and the dynamic is much simpler (as there is only 3 dimensions).

**Figure 6:**
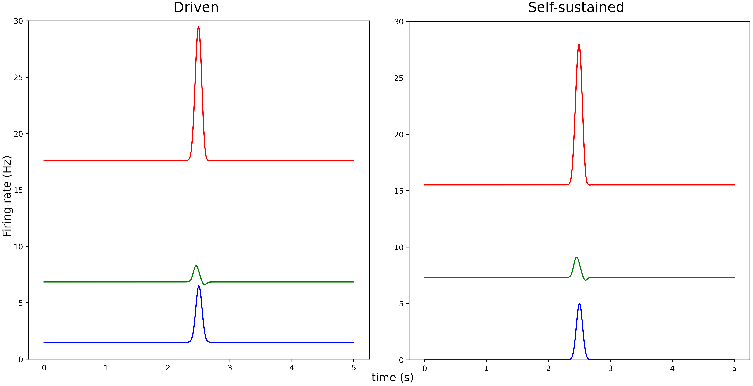
Response to a perturbation centered at 2.5s. Green represent the excitatory population, red is in inhibitory one, and blue is the external drive. Left part is for a driven mean-field, right part for a self-sustained one.

Table.6 is identical to Table.5, appart from the absence of different noise realizations, as no noise were used in the mean-field. It presents a similar result to Table.5 : self-sustained mean-field do indeed appear to have a higher response than driven ones.

**Table 6:** Response of the Mean-Field with different external currents (in lines) for 2 Driven and 2 Self-sustained networks and for the two neuronal populations (in columns) : excitatory and inhibitory. In each cells, we have *Resp*_*abs*_ followed by *Resp*_*perc*_

| Driven | Exc. FR 1 | Exc. FR 2 | Inh. FR 1 | Inh. FR 2 |
| --- | --- | --- | --- | --- |
| Input = 3Hz | 0.751,11.1% | 1.348,13.0% | 6.627,37.6% | 66.319,28.7% |
| Input = 5Hz | 0.869,12.7% | 1.745,16.8% | 10.228,58.1% | 9.694,44.0% |
| Self-sustained |  |  |  |  |
| Input = 3Hz | 0.994,13.6% | 1.771,16.8% | 7.160,46.2% | 7.026,35.4% |
| Input = 5Hz | 1.132,15.5% | 2.230,21.1% | 10.814,69.7% | 10.509,52.9% |

It appears that, both for the AdEx networks and mean-fields, self-sustained ones (which had higher LE) have a higher response than driven ones.

## 4 Discussion

In this work, we analyzed two types of neural networks, self-sustained networks and driven metworks. We used the Lyapunov exponent, a dynamical system tools, to yield insight on the nature of their dynamics. We also investigated the responsiveness of such networks, and found that the self-sustained networks show a tendency to be more responsive than driven networks. This difference can be captured by mean-field models. We discuss here these results and their implications.

One main finding is that the FLE is different for the two types of AdEx networks. It must be noted here that we did not technically computed the FLE of the network, as the AdEx networks have 40 000 dimensions (and the HH networks have even more). As it would have been difficult to use algorithms in such a high-dimensional space, partly due to the curse of dimensionality (Köppen, 2000), we restricted our use to a system made from the average values of the FRs and the Adaptation (or the other variables for HH). It is interesting to note that the FRs are directly correlated to a change of activity of the network, while the link between it and the Vm is more tenuous (due to the reset after the spike) and high or low values of Vm are hard to link with the global activity of the network. This could explain why we obtained interesting results with the FR but not with the Vm.

The computation of the FLE (as seen in Table.1) indicates that both types of network are slightly chaotic (and could be way more chaotic under 40 000 dimensions instead of 3), as they exhibit weakly positive FLE. But our main result is that Self-sustained networks had a FLE roughly an order of magnitude higher than Driven networks. Of course, a major difference between Driven and Self-sustained systems, by definition, is that one of them has a constant external drive applied to it while the other does not. It seems obvious that this potential added drive would influence the dynamics one way or another, and that this influence would be found in the system made from the average values as well. It could have very well been that we observed only the difference due to the drive, that would have caused a decrease in the LE somehow. This is why we did simulations with the same Self-sustained networks as before, but with an external drive similar to the one we used for Driven networks. We then computed their FLE, as seen in Table.2. If the difference between Driven and Self-sustained was due to the drive, then similarly as before, the FLE should be lower, but this is not the case : they are roughly of the same values, and higher drives do not seem to increase or decrease much of the FLE. This means that the difference we observed is not due to an added external drive, but due to the intrinsic dynamics and specific sets of parameters that create the different types of network.

Furthermore, different set of parameters were used to obtain the four networks, two Driven and two Self-sustained, and despite having only those parameters from the single neurons model changed, differences between their FLE were observed between different types of networks, but not within the same type (or at least, not to the same extent).

In other words, two networks that look similar if one only look at their activity through the FR, are actually intrinsically different. The difference in the FLE means Self-sustained networks are more chaotic than Driven ones. This means they both behave differently, which can have an impact on how they interact with other systems are tool to measure them, or how they would change when parameters are changed.

As can be seen in Table.3, the result we just described is only valid for the AdEx model.

There are a few possible reasons explaining why we see a difference between the two types for AdEx but not for HH networks : it could be that the difference also exists but is not capture by our method of reconstructing a system, or that the difference is unique to AdEx networks, or that the difference uniquely does not work in HH networks but would work with, said, other integrate and fire models.

We want to emphasize that even if the result we show was only true for AdEx (which is not obvious yet), that would already be useful for studying AdEx itself, obviously, but also to study the dynamical differences between the networks created from AdEx and other models.

A second main result concerns the responsiveness of the two different types of networks. We found that self-sustained networks generally display a higher responsiveness compared to driven networks (Table.5). However, this does not mean all Self-sustained networks would have a stronger response than Driven ones : there is a huge difference between various networks of the same type, and numerous parameters could influence the response to a perturbation. Here we merely show a tendency for a few specific examples. Future studies should scan the parameter space of such models in more detail to determine if this feature is universal.

It is important to note that the link between LE and responsiveness was shown in another study (di Volo & Destexhe, 2021), but the responsiveness was more related to the second Lyapunov exponent of the system. Interestingly, we found a similar correlation between responsiveness and second exponent in the mean-field model (Table.4 and Table.6). Such a relation between the LE spectrum and responsiveness, or more generally between stability and responsiveness, is also an interesting direction to explore in future work.

## Acknowledgments

Research supported by the CNRS, Agence Nationale de la Recherche (ShootingStar project, AD), The European Union (H2020-945539, AD), and a PhD fellowship from the Initiativas Foundation (JB).

## Supplementary

The code showing the algorithms and simulations can be find at https://github.com/JuBoute/lyapunov neural network

